## Supplementary material for "Tiered Sympathetic Control of Cardiac Function Revealed by Viral Tracing and Single Cell Transcriptome Profiling": Table S1

### Supplementary Table 1

| Sample | Batch | Sequencing Depth  (UMIs/Cell) | Sequencing Saturation |
| --- | --- | --- | --- |
| Mouse-2-1_WTLM | 2 | 5,558 | 60.9% |
| Mouse-2-2_WTLM | 2 | 6,241 | 68.3% |
| Mouse-2-3_WTLM | 2 | 3,818 | 83.1% |
| Mouse-2-4_WTLM | 2 | 5,627 | 59.8% |
| Mouse-2-6_WTLM | 2 | 4,953 | 60.4% |
| Mouse-1-3_WTLM | 1 | 5,599 | 88.5% |
| Mouse-1-6_WTLM | 1 | 4,441 | 87.8% |
| Mouse-1-7_WTLM | 1 | 5,157 | 81.2% |
| Mouse_3-1_DCM | 3 | 4,883 | 97.1% |
| Mouse_3-2_DCM | 3 | 3,230 | 97.8% |
| Mouse_3-3_WTLM | 3 | 5,486 | 79.8% |
| Mouse_3-4_WTLM | 3 | 5,641 | 90.5% |
| Mouse_3-5_WTLM | 3 | 5,841 | 94.1% |
